# Detecting apoptosis via DODO

**DOI:** 10.1101/2022.05.14.491923

**Authors:** Ziheng Zhang, Zhe Sun, Ji-Long Liu

## Abstract

The real-time detection of intracellular biological processes by coded sensors has broad application prospects. Here we develop a degron based modular reporting system: the <u>D</u>evice <u>o</u>f <u>D</u>eath <u>O</u>peration (DODO), which can be used to detect a series of biological processes. The DODO system consists of “reporter”, “sensor” and degron. After protease activation and cleavage, the degron will be released from the fluorescent protein and eventually lead to the stabilization of the fluorescent protein. By replacing different “sensors” and “reporters”, a series of biological processes can be reported through different signals. The system can effectively report the existence of TEV. To prove this concept, we successfully apply the DODO system to report apoptosis. In addition, the reporter based on degron will help to design protease reporters other than caspase.

## INTRODUCTION

The techniques of detecting intracellular signals play important roles in biological research, especially for those key cellular molecules representing basic biological processes. Informative detections include the detection of the expression of some key genes, the translocation of some proteins and the activation of zymogens.

To gain insight into the physiological state of cells, the common practice is to determine the expression, location and modification of key biomarkers through a series of biochemical experiments. In the past decades, people have developed a series of detection methods for different physiological states and signal molecules. In particular, biological coding reporting systems, such as coupling conformationally sensitive cpEGFP with human dopamine receptors to detect dopamine^1^. Similar methods include calcium sensors and acetylcholine sensors^2–3^. In general, these methods are tailored to the characteristics of different targets.

Signals of many intracellular events are transmitted through the activation of zymogens. A classic case based on zymogen activation is apoptosis, which is performed by caspase^4–5^. Apoptosis has always been the focus of attention because it is related to developmental events, cancer treatment and many diseases^5–6^. In the past many years, people have developed a series of methods to detect apoptosis. Mainly by detecting the activation of key signaling molecules, such as caspase3. The activation of zymogen plays a fundamental role in many intracellular pathways, especially the report of zymogen activation is very important.

Recently, it has been reported that a series of degrons are mainly used to artificially regulate protein degradation^7–9^. A famous example is AID, a powerful tool developed from auxin receptors in in plants^7^. F-box transport inhibitor reaction 1 (TIR1) is a receptor for auxin. By binding to auxin, chimeric E3 ligase (SCF-TIR1) recruits and ubiquitinates the degron^10–11^. This is a natural and rapid method of regulating substrate proteins. Fortunately, the orthologs of TIR1 is only found in plants, which provides the possibility to apply it to other species. Based on this principle, AID has developed into a powerful technology for conditional and rapid degradation of target proteins. By exogenously expressing TIR1 and adding auxin or other analogues to the medium, such as IAA (indole-3-acetic acid) and NAA (1-naphthylacetic acid), the degron labeled protein will degrade rapidly^7^, ^12^. Driven by gene knock-in technology based on CRISPR / Cas9, it is widely used for rapid degradation of endogenous proteins^12–13^.

In addition, many proteins have natural unstable motifs at the N-terminal or C-terminal, which may be composed of a few amino acids. Large scale N-segment screening or C-terminal screening can be used to characterize these unstable motifs^14–15^. These results have been confirmed by some natural proteins and can be used to develop a series of tools ^16–17^. This is a suitable element for muting the reporting signal in the control unit. In this report, we introduce the concept of modularity, which combines a reporter, an inducer and a degron into a complete reporting system, which we name as “the <u>D</u>evice <u>o</u>f <u>D</u>eath <u>O</u>peration (DODO)”. DODO can be used to report the activity of target enzymes in living cells and be applied for indentifying apoptosis in vivo. Moverover, people can change these elements according to the needs of different purposes.

## RESULTS

### Strategies for developing DODO

DODO, a modular self-degradation reporting system, consists of three components, a reporter, an inductor and a degron. Among them, the reporter is used to report signals; the inductor is used to monitor specific signals and the degron is used to trigger its own degradation. When there is no target signal in the cell, it will trigger the degradation of reporter gene and make the cell negative. Otherwise, the sensor will cut off the degron, stabilize the reporter and accumulate in the cell, and finally be detected (Fig. 1a). In this system, the reporter can be determined according to needs, such as fluorescent protein, luciferase or some labels; The inductor depends on the target signal. For example, the apoptosis inductor we used in this study is caspase3 cleavage sequence (DEVDG); For degron, we choose C-degron (RRRG), which has strong degradability^16^.

**Fig. 1.**
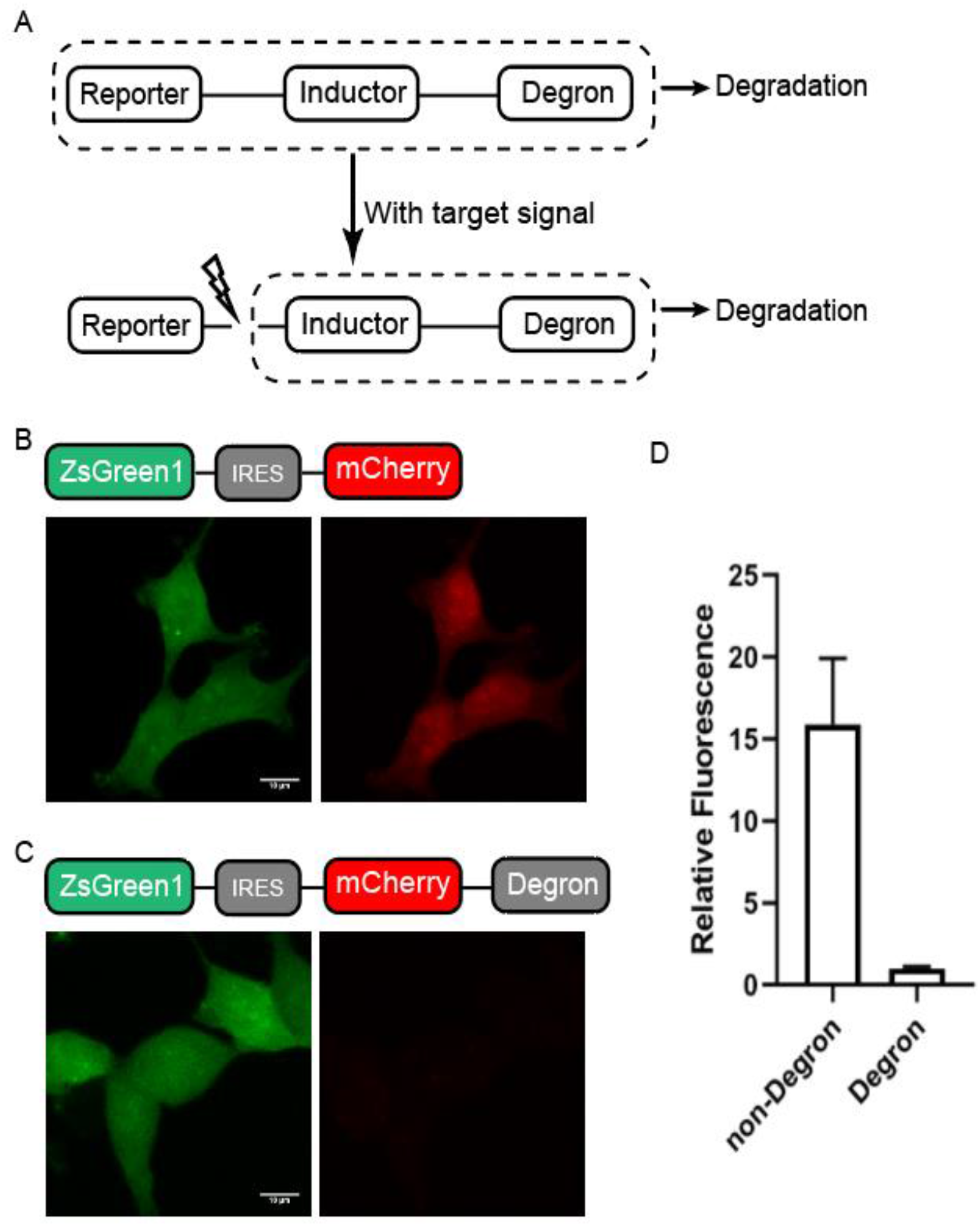
Design of DODO. A. Design and working principle of self-degradation reporting system. B. The stable expression of EGFP in cells and the co-expression of mCherry mediated by IRES under the same promoter. C. Degron and EGFP labeled mCherry were stably expressed in the same promoter. D. The relative fluorescence intensity of mCherry with and without Degron.

Background noise is usually an important factor that makes it impossible to accurately determine the authenticity of the target signal. Therefore, we first determined the degradation ability of degron. We use fluorescent protein to achieve this. By adding degron to the C-terminal of fluorescent protein to trigger the degradation of fluorescent protein, we can judge its degradation ability. The specific method is to construct green fluorescent protein and IRES linked red fluorescent protein under the same promoter, add or not add degron at the C end of red fluorescent protein, and test the degradation ability of degron (Fig. 1 b, 1c). After imaging the red fluorescent protein signal under the same confocal microscope parameters, it can be seen that the red fluorescent protein with degron has almost no red signal compared with the signal without degron (Fig. 1b and 1c). By comparing the red fluorescence intensity under the two conditions, it can be determined that degron can degrade about 95% of the target protein.

### DODO responds to target signals

To test whether the DODO system can respond to specific signals, we introduce a TEV cleavage site between the reporter and degron as an inductor, and detect whether the presence of TEV can be reported. TEV has categoric sites and high specificity, and has been widely used in a series of applications including protein purification. The presence of TEV was detected and reported by introducing the cleavage sequence of TEV between the reporter and degron (Fig. 2a).

**Fig. 2.**
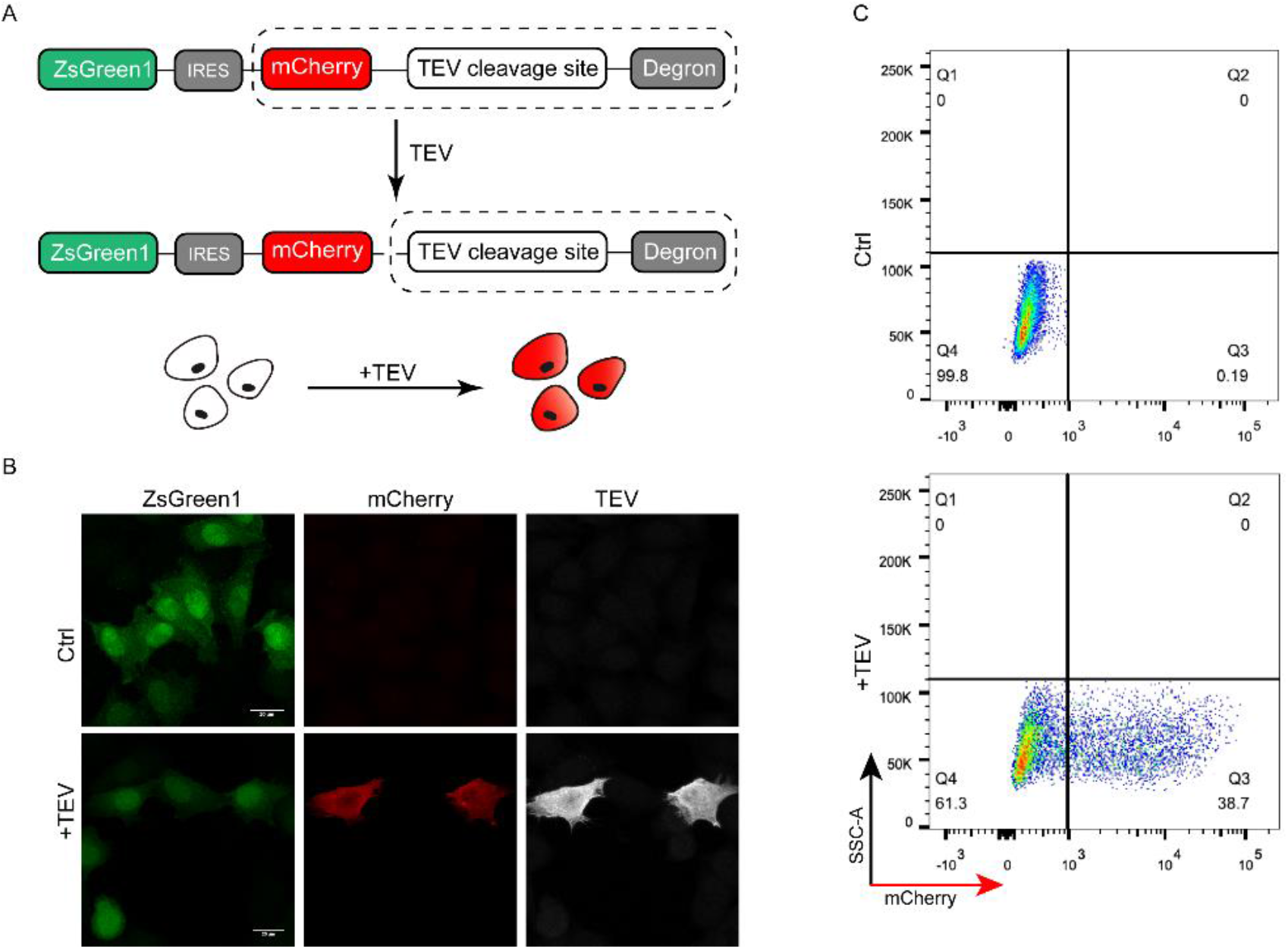
Validation of DODO by TEV and TEV cleavage sequence. A. The pattern diagram shows the introduction of TEV cleavage sequence between mCherry and degron to detect whether DODO can report the presence of TEV signal. B Fluorescence of cells stably expressing EGFP-IRES-mCherry-TEV cleavage sequence-degron with and without TEV transfection. C. Thirty-six hours after transfection of pCMV-TEV-myc, the fluorescence signal of the cells was analyzed by flow cytometry. D. Relative fluorescence intensity of the cells with or without TEV.

In order to ensure the consistency of data, we constructed a stable monoclonal stable cell line to verify whether the system can respond to TEV. By transiently transfecting pCMV-TEV-myc into a stable cell line, the successfully transfected cells showed mCherry positive (Fig. 2b). Further flow cytometric detection of cells after pCMV-TEV-myc transduction showed that the cells could respond to TEV and were mCherry positive (Fig. 2C).

### DODO detects apoptosis in 2D culture

The above experiments have confirmed that DODO can effectively respond to the emergence of target signals. We further applied DODO to the report of specific signals in cells. There are many kinds of signal molecules with protease activity in cells, among which caspase family is the most typical and well-known. We applied this modular self-degradation reporting system to report apoptosis by introducing the cleavage site of caspase3 between mCherry and degron. By reporting the activation of caspase3, it was shown whether the cells had apoptosis (Fig. 3a).

**Fig. 3.**
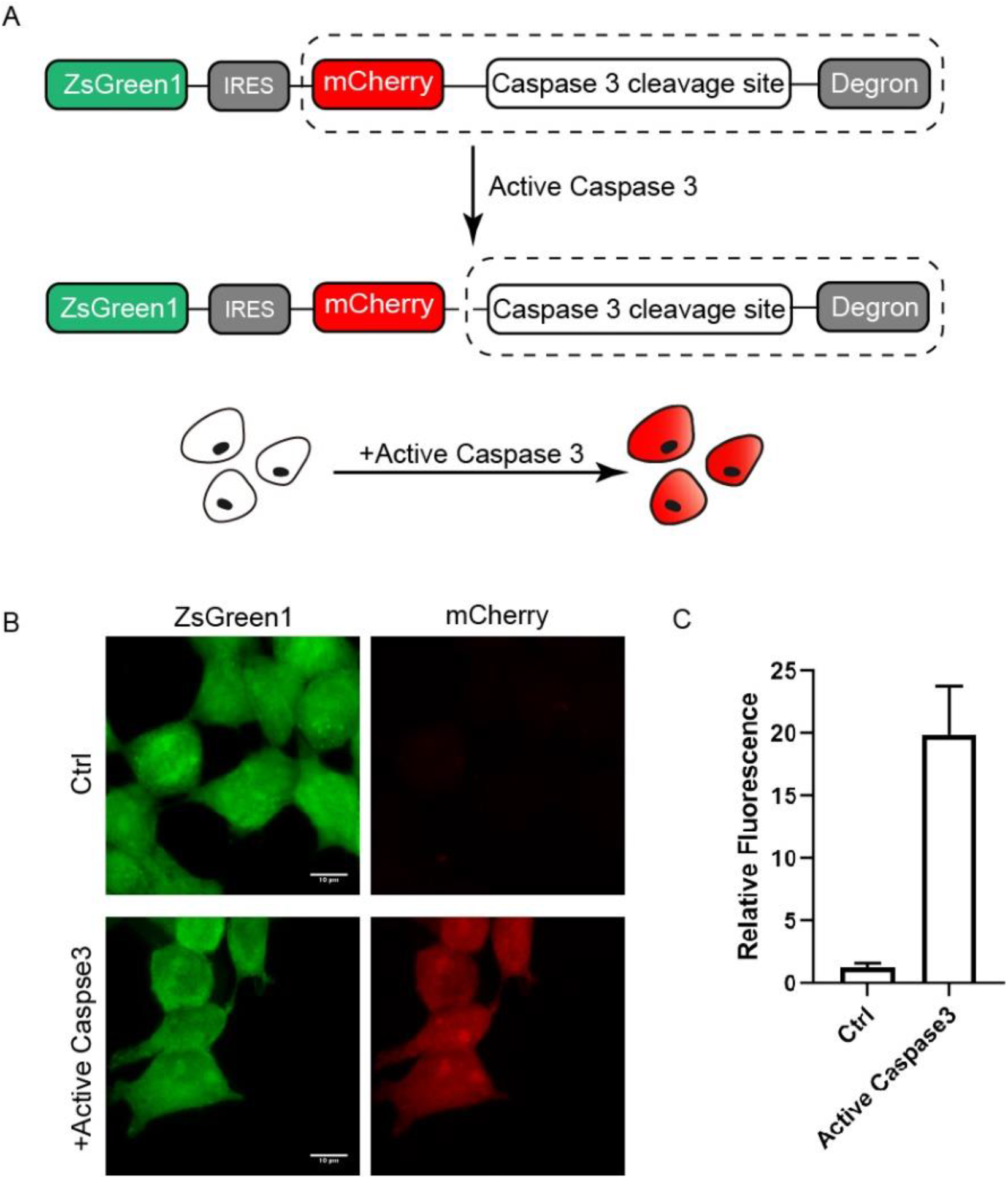
DODO responds to activated caspase3. A. The pattern diagram shows the introduction of caspse3 cleavage sequence between mCherry and degron to report the activation of caspase3. B. Fluorescence of cells after transfection of pCMV-active caspase3. D. Relative fluorescence intensity of cells transfected or not transfected with active caspase3

We first simulated the activation of caspase3 in cells by transient transfection of active caspase3, and further determined whether the reporting system could respond to activated caspase3. After expressing exogenous active caspase3, the cells were mCherry positive. In contrast, the intensity of mCherry signal increased about 20 times (Fig. 3b and c).

### DODO detects apoptosis in 3D culture

The above experiments show that DODO can be used to report the target signal molecules with protease activity in 2-D cultured cells. In order to further apply the system to report apoptosis under physiological conditions, we further tested the system in MCF-10A cells. MCF-10A has the characteristics of forming cell clusters in 3D culture in vitro. Under 3D culture, a single MCF-10A cell will gradually grow from a single cell to a cell cluster. Over time, the cells located in the cell cluster will undergo apoptosis and finally form a cavity (Fig. 4a).

**Fig. 4.**
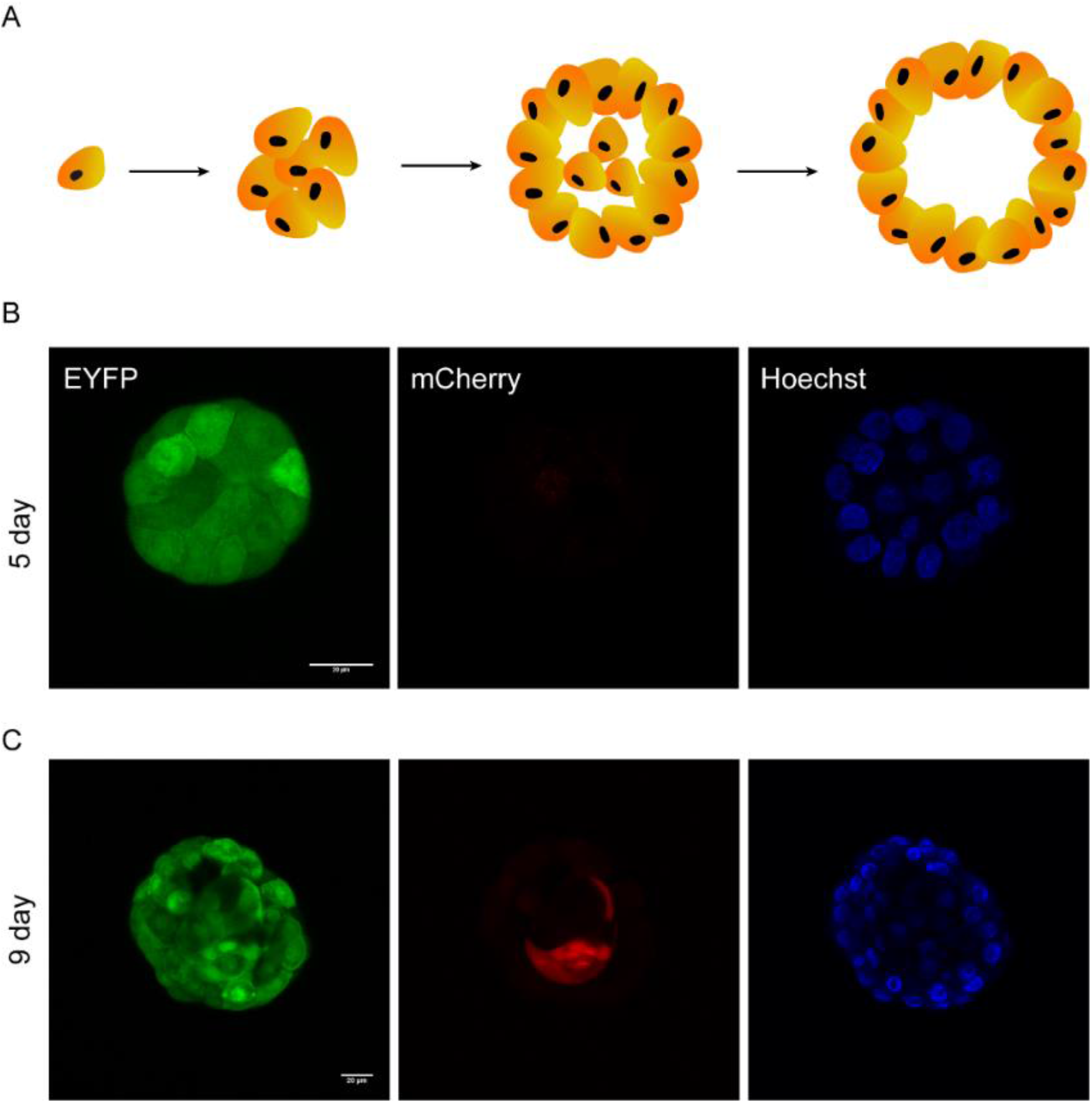
DODO determines the apoptosis of MCF-10A during metamorphic development in 3D culture. A. In the schematic diagram of the metamorphic development process of MCF-10A in 3D culture, the cells in the center of the cell cluster will gradually undergo apoptosis and eventually form a cavity. B. The cell cluster was proliferative without apoptosis, and all cells were mCherry negative. D. The cell cluster was in the state of apoptosis, and the central cells began to apoptosis, labeled as mCherry positive.

This process is mainly divided into two stages. One is the proliferation stage, which starts from about 1 ~ 8 days. At this stage, cells mainly divide and proliferate to form cell clusters. The second stage is apoptosis (about 7 ~ 13 days). Due to the limitation of malnutrition, the cells located in the cell cluster begin to apoptosis. The 3D culture of MCF-10A is a good model for reporting apoptosis^18^ and for testing the sensitivity of DODO.

We used lentivirus to construct MCF-10A cells expressing mCherry-caspase3_cleavage_site-IRES-EYFP stably. Matrigel was used to support 3D culture of MCF-10A cells. We tried to obtain images of different stages, including proliferation stage, to test whether the apoptosis reporting system will incorrectly report cells in proliferation stage and whether it can report apoptotic cells in apoptosis stage.

We took images of MCF-10A cell clusters on day 6 of 3D culture (Fig. 4b). No matter the cells located in the outer layer or the cells located in the inner layer, the cell clusters do not show mCherry signal. This shows that degron can maintain the ability to degrade the target protein in this environment, and it also shows that the reporting system will not misreport in the stage of cell proliferation.

We also imaged MCF-10A cells in the apoptotic state (Fig. 4c). A specific internal cell showed mCherry positive, indicating that caspase3 of these cells was in the caspase3 activated state. The above data show that the reporting system can effectively report the apoptosis of MCF-10A cells in 3D culture.

## DISCUSSION

The codable tools for reporting cellular biological processes are of great significance for people to identify the physiological state of cells. In particular, modular tools can provide researchers with a variety of options and reduce application restrictions.

In this report, we describe a new degron-based modular reporting system, DODO, for the effective detection of a series of biological processes. DODO is mainly used for a series of intracellular signal proteins with protease activity. By introducing the target sequence of specific signal protein with protease activity between reporter and degron, the report of specific intracellular signal and cell physiological state can be realized.

A previous report described a GFP based fluorescent protease reporter gene FlipGFP, which reverses a beta strand of the GFP. After protease activation and cleavage, the beta strand recovery leads to the reconstruction of GFP and fluorescence^19^. Recently, FlipGFP has been used as a tool for high-throughput identification of inhibitors of 3CL^pro^ of SARS-CoV-2^20^. This provides a new strategy for reporting target signals, but there are still some limitations. Just like FlipGFP, the reporter is fixed, which leads to the unity of its detection form (although there are also red fluorescent versions). People cannot change the reporter according to their own needs. DODO, as a modular reporting system, provides researchers with wider applicability, which can be assembled into appropriate tools according to their needs and replaced simply.

In conclusion, we develop DODO as a codable and degron-based modular protease reporting system in this study. By changing the components of the system, different protease signals can be reported in a variety of reporting ways.

## MATERIALS AND METHODS

### Cell culture

For HEK293T, cells were grown in DMEM (Hyclone, SH30022.01) containing 10% FBS (Biological Industries, 04-001-1A) in a humidified incubator at 37 °C with 5% CO2. For MCF-10A, cells were grown in Growth Medium, consists of DMEM/F12 (Corning, 10-092-cvr) with 10% House Serum (Biological Industries, 04-124-1A), 20ng/ml EGF (Peprotech, AF-100-15-100), 0.5 mg/ml Hydrocortisone (Selleck, S1696), 100 ng/ml Cholera Toxin (APExBIO, B8326) and 10 ug/ml Insulin (Medchemexpress, HY-P1156) in a humidified incubator at 37 °C with 5% CO2.

### Plasmid construction

All plasmids were created by standard molecular biology techniques and confirmed by exhaustively sequencing the cloned fragments. In particular-for pLV-ZsGreen-IRES-mCherry-TEV-degron, we introduced the TEV cleavage sequence (5’-GAGAACCTGTACTTCCAGAGC-3’, Amino acid sequence ENLYFOS) between the C-terminal of mCherry and degron by ClonExpress Ultra One Step Cloning Kit (Vazyme, C115-01). For the plasmid to report apoptosis, we replaced the TEV cleavage sequence with the caspase3 cleavage sequence (5’-GGAGACGAGGTGGACGGC-3’, Amino acid sequence DEVDG).

### Lentivirus packaging and stable cell line construction

All plasmids were prepared using E.Z.N.A.® Endo-free Plasmid DNA Mini Kit (Omega, D6950-01B) and transfected with Lipofectamine 2000 (Thermo, cat. #11668019). 293T cells were cultured in 6cm dish. Co-transfected with packaging vector psPAX2 (Addgene#12260), envelope vector pMD2.G (Addgene#12259) and transfer vector pLV-ZsGreen-IRES-mCherry-TEV-degron or pLV-ZsGreen-IRES-mCherry-Casp3-degron into 293T cells in a ratio of 3:1:4 (totally 5μg for 6cm dish) when the cell density is about 70%. Forty-eight hours after transfection, the viral supernatant is collected and filter supernatant through a 0.45 μm PES filter. 1 ml of virus supernatant were used for cell infection. Seventy-two hours after infection with lentivirus, the cells with EGFP-positive and mCherry-negative were collected by flow sorting.

### Immunofluorescence

For immunofluorescence experiment, the 293T-ZsGreen-IRES-mCherry-TEV-degron were transfected with pcDNA3.1-TEV-myc. Twenty-four hours after transfection, cells were washed with PBS, and fixed through 4% PFA at room temperature 15min. 5% BSA was used for blocking, and incubated with anti-myc (Santa Cruz, sc-40) antibody overnight at 4°C. After washed with PBST (PBS with 0.1% TritonX-100), cells were incubated with secondary antibody (anti-mouse cy5, 715-175-151) at room temperature for 1 hour, after mounting the slides were used for image.

### 3-Dimensional culture of MCF-10A

For of MCF-10A, before 3Dimensional culture, MCF-10A were grown in Growth Medium, consists of DMEM/F12(Corning,10-092-cvr) with 10% House Serum (Biological Industries, 04-124-1A), 20ng/ml EGF (Peprotech, AF-100-15-100), 0.5 mg/ml Hydrocortisone (Selleck, S1696), 100 ng/ml Cholera Toxin (APExBIO, B8326) and 10 ug/ml Insulin (Medchemexpress, HY-P1156) in a humidified incubator at 37 °C with 5% CO2. For 3D culture, the cells are digested by 0.05% Trypsin-EDTA, and resuspend in 3-5 mL Resuspension Medium (DMEM/F12 with 10% House Serum). Centrifuging to harvest the cells, and resuspend in 1 mL of Assay Medium (DMEM/F12 with 10% House Serum, 0.5 mg/ml Hydrocortisone, 100 ng/ml Cholera Toxin and 10 ug/ml Insulin). Filter with cell strainer and count cells, make the cell concentration as 12,500 cells/ml in Assay Medium with 2% Matrigel (Corning, 354230) and 5ng/ml EGF. Plate 400 μL of this mixture on top of the solidified Matrigel in each well of the chamber slide. This corresponds to a final concentration of 5,000 cells/well in Assay medium containing 2% Matrigel and 5 ng/ml EGF. Take the cells to grow in incubator at 37 °C with 5% CO2. The cells should be refed with Assay Medium containing 2% Matrigel and 5 ng/ml EGF every two days. Cells can be used for imaging at the desired timepoint.

## ACKNOWLEDGMENTS

This work was supported by grants from the Ministry of Science and Technology of the People’s Republic of China (grant no. 2021YFA0804701-4), the National Natural Science Foundation of China (grant nos. 31771490 and 81972632), the Shanghai Science and Technology Commission (20JC1410500). We thank the Molecular Imaging Core Facility (MICF) at School of Life Science and Technology, ShanghaiTech University for providing technical support.

